# Influence of the broadly neutralizing antibody VRC01 on HIV breakthrough virus populations in antibody-mediated prevention trials

**DOI:** 10.1101/2025.07.01.659979

**Authors:** Carolyn Williamson, Chivonne Moodley, Craig A. Magaret, Elena E. Giorgi, Morgane Rolland, Dylan H. Westfall, Anna Yssel, Wenjie Deng, Raabya Rossenkhan, Nonhlanhla N. Mkhize, Lennie Chen, Hong Zhao, Tanmoy Bhattacharya, Alec Pankow, Ben Murrell, Talita York, Asanda Gwashu-Nyangiwe, Nonkululeko Ndabambi, Ruwayhida Thebus, Paula Cohen, Bronwen Lambson, Haajira Kaldine, Sinethemba Bhebhe, Michal Juraska, Hongjun Bai, Allan C. deCamp, James Ludwig, Cindy Molitor, Nicolas Beaume, David Matten, Yunda Huang, Lily Zhang, Daniel B. Reeves, Bryan Mayer, Shelly T. Karuna, John A. Hural, Lynn Morris, David Montefiori, Roger E. Bumgarner, Penny L Moore, Paul T. Edlefsen, Srilatha Edupuganti, Nyaradzo Mgodi, M. Juliana McElrath, Myron S. Cohen, Lawrence Corey, Peter B. Gilbert, James I. Mullins

**Author notes:** **Correspondence:** Carolyn Williamson, Division of Medical Virology, Institute of Infectious Disease and Molecular Medicine Faculty of Health Sciences, University of Cape Town and National Health Laboratory Service, South Africa, James I. Mullins, Department of Microbiology, University of Washington, Seattle, WA, USA. Department of Microbiology, The Icahn School of Medicine at Mount Sinai, New York, NY 10029, USA. These authors contributed equally to this work (Chivonne Moodley, Craig A. Magaret). These authors contributed equally to this work (Carolyn Williamson, James I. Mullins).

## Abstract

In the HIV antibody mediated prevention (AMP) trials, the broadly neutralizing antibody VRC01 demonstrated protective efficacy against new diagnoses with susceptible HIV strains. To understand how VRC01 shaped breakthrough infections, we performed deep sequencing on 172 participants in the placebo and treatment arms, generating 63,444 *gag-Δpol* (2.5 kb) and 53,088 *rev-env-Δnef* (3 kb) sequences. Sequences were classified into transmitted founder lineages (TFLs), and infections with multiple distinct lineages were determined. Multilineage infections were detected in ∼38% of participants in both the African (HVTN 703/HPTN 081) and Americas/Europe (HVTN 704/HPTN 085) cohorts, regardless of placebo or treatment group, or cohort. The high levels of multilineage infections could be attributed to minor lineages (<5% abundance) identified in 20% of participants. Infection with VRC01 discordant viruses (IC80s >3-fold different) was observed in 40% of multilineage infections, with a trend toward greater intra-host neutralization differences with increasing VRC01 dose (Jonckheere-Terpstra test, p=0.072). In six VRC01 treated participants who acquired both sensitive (IC80<1µg/ml) and resistant viruses (IC80>3µg/ml), the sensitive lineages declined over time. Recombination was pervasive, observed in 63% of multilineage infections at the time of HIV diagnosis. In one treated participant infected with VRC01 discordant lineages, recombinant viruses preferentially inherited the resistance mutation (binomial p=0.004). In conclusion, our in-depth analysis of breakthrough viruses in the AMP trials revealed a high frequency of multilineage infections, including infections with viruses with different VRC01 sensitivities. This analysis also highlights the role of recombination in shaping intra-host viral evolution and facilitating escape from VRC01.

**Significance:** This work advances our understanding of the diversity of initial viral infection and evolution and in vivo activity of broadly neutralizing antibodies (bNAbs). Deep sequencing revealed ∼38% of HIV acquisitions with multiple transmitted founder lineages, higher than previous studies. These occurrences were similar in the placebo and VRC01 groups. Viral recombination among post-acquisition variants was common under antibody selection and appeared to favor resistant sequences in the treatment group. These data suggest as with single antiviral therapy, passive and active immunization of bNAbs should be directed at multiple antigen targets of HIV-1.

## INTRODUCTION

HIV remains a significant global public health concern, and utilizing broadly neutralizing antibodies (bNAbs) for prevention is a crucial component of the strategy to combat its spread. The only efficacy studies assessing this concept, the antibody mediated prevention (AMP) trials, evaluated the effectiveness of the bNAb VRC01 (10 doses given every 8 weeks at either 10 mg/kg or 30 mg/kg) to prevent HIV acquisition. These trials were conducted in women in sub-Saharan Africa (HVTN 703/HPTN 081), and in cismen and transgender persons who have sex with men in the Americas and Europe (HVTN 704/HPTN 085) (1). The AMP trials showed that although VRC01 had only low overall prevention efficacy (PE) against overall HIV diagnosis, it had 75% PE against viruses sensitive to VRC01 (IC80 < 1 µg/ml) (1). Viruses acquired in the VRC01 groups were more resistant to neutralization by the antibody than those in the placebo group (geometric mean IC80 of 8.4 µg/ml compared to 3.5 µg/ml, respectively) and exhibited greater divergence in the VRC01 epitope (1-3). Together, these findings underscore the complexity of the interaction between bNAbs and HIV and emphasizes the importance of characterizing the genotypic and neutralization characteristics of the infecting virus population to inform biomedical prevention strategies.

In people living with HIV (PLWH), the virus persists as genetically related but evolving and diversifying genome populations, also known as “quasispecies”, although only one to a few virions typically establish clinical infection (4-9). Because we cannot know exactly which specific variants within a closely related population crossed the mucosal/blood barrier and proliferated in the ensuing infection, in this paper we refer to the viral populations that emerge after acquisition as transmitted founder lineages (TFL). Following transmission, there is a selection bias for fitter viruses (9) which preferentially use the CCR5 co-receptor and tend to be more interferon-alpha resistant (10-13). In contrast, factors that disrupt the mucosal barrier, such as other sexually transmitted infections or genital inflammation, reduce this selection bias, resulting in a greater number of viruses establishing infection, and particularly viral strains with lower infectivity (9, 14, 15).

A meta-analysis of 70 studies estimated that 25% of individuals acquire multiple founder viruses, with higher rates in men who have sex with men (∼30%) than in women (∼21%)(16). Traditional sequencing approaches had limited sensitivity, but recent advances in long-read deep sequencing now allows more comprehensive analysis of viral quasispecies (17). Using this method, we generated over 100,000 *gag-Δpol* (GP) (2.5 kb) and *rev-env-Δnef* (REN) sequences from the AMP trials. Using high-resolution sequencing, we identified ∼38% multilineage infections – a higher rate than reported in lower-sensitivity studies (Baxter et al., 2023). The prevalence remained consistent across both placebo and treatment arms (low and high dose VRC01 groups) and both cohorts. Of those with multilineage infection, 40% were infected with viruses with differing VRC01 sensitivities (difference in IC80 > 3-fold). In the treatment group, the frequency of VRC01 sensitive lineages decreased over time, suggesting post-acquisition suppression of sensitive lineages, and selection of resistant lineages. Recombination was pervasive in very early infection, and in one participant, recombinant viruses preferentially inherited the VRC01-resistant mutation, highlighting the role of recombination in early viral adaptation under selective antibody pressure.

## RESULTS

### Description of participants and data generated

AMP trial participants were infused for 10 visits, every 8 weeks, with either 10 mg/kg VRC01, 30 mg/kg VRC01, or saline (placebo). Individuals were included if they were diagnosed with HIV by the week 80 visit (primary endpoints) **(Table 1)**. Those who acquired HIV after the week 80 visit were excluded due to declining VRC01 levels at the time of diagnosis. Individuals were tested for HIV acquisition every 4 weeks, and sequencing was performedon the first HIV RNA positive sample. In 159 individuals, at least one additional sample was sequenced - collected within a median of 13 days (IQR 6-21) after the first sequencing time point.

**Table 1:** Genotypic and neutralization characteristics of viruses in participants with HIV diagnosis by week 80 visit (primary endpoint).

|  | HVTN 703/HPTN 081 (Africa Trial) |  |  |  | HVTN 704/HPTN 085 (Americas Trial) |  |  |  |
| --- | --- | --- | --- | --- | --- | --- | --- | --- |
|  | Placebo | VRC01 10 mg/kg | VRC01 30 mg/kg | Total | Placebo | VRC01 10 mg/kg | VRC01 30 mg/kg | Total |
| No. of primary endpoints* | 29 | 28 | 17 | 74 | 38 | 32 | 28 | 98 |
| No. of REN multilineage infections | 11 | 8 | 9 | 28 | 13 | 13 | 10 | 36 |
| No. of GP multilineage infections | 11 | 7 | 8 | 26 | 14 | 12 | 6 | 32 |
| No. of multilineage infections (GP and/or REN) (%) | 11 (38%) | 8 (29%) | 9 (53%) | 28 (38%) | 15 (39%) | 13 (41%) | 10 (36%) | 38 (39%) |
| No of multilineage infections with IC80 data on >1 lineage | 6 | 6 | 5 | 17 | 9 | 8 | 6 | 23 |
| No. of VRC01 discordant multilineage infections** | 1 (17%) | 3 (50%) | 2 (40%) | 6 (35%) | 3 (33%) | 3 (38%) | 4 (67%) | 11 (48%) |
| No. of REN sequences first timepoint: mean (range) | 134 (6-493) | 115 (4-550) | 143 (6-625) | 129 (4-625) | 175 (4-840) | 148 (2-635) | 117 (1-381) | 150 (1-840) |
| No. of GP sequences first timepoint: mean (range) | 117 (6-474) | 101 (3-374) | 141 (11-981) | 116 (3-981) | 159 (26-377) | 180 (1-1055) | 113 (6-235) | 151 (1-1055) |
| No. of REN sequences second time point: mean (range) | 123 (3-353) | 168 (14-546) | 223 (30-633) | 163 (6-633) | 149 (38- 83) | 224 (2-1128) | 164 (31-421) | 178 (2-1128) |
| No. of GP sequences second time point: mean (range) | 167 (9-526) | 211 (22-701) | 227 (10-814) | 199 (9-814) | 145 (24-396) | 150 (4-455) | 107 (1-267) | 136 (4-455) |

\*Primary endpoints include those participants with available first RNA+ sample sequencing data. The total endpoints in the Africa trial were 76, and two were excluded.
\*\*Pseudoviruses where at least one pair were >3-fold difference in IC80 titres. REN: rev-vpu-env-Δnef. GP: gag-Δpol.

Long-read, deep sequencing data were generated from 172 individuals who acquired HIV in AMP: 74 from the HVTN 703/HPTN 081 Africa trial and 98 from the HVTN 704/HPTN 085 Americas trial **(Table 1)**. Two viral genomic regions were sequenced at each time point: *gag-Δpol* (GP) (2.5 kb) and *rev-env-Δnef* (REN) (3 kb), resulting in a total of 57,798 GP and 53,088 REN sequences, with a mean of 162 GP and 147 REN sequences generated per participant per timepoint. Neutralization data were generated using gp160 env plasmids representative of TFLs from 162 individuals: 72 individuals from the Africa trial, and 90 individuals from the Americas trial (2).

### Lineage classification

Using data from both time points, HIV sequence populations were classified into lineages, referred to as Transmitted Founder Lineage (TFL), with each lineage assumed to represent a distinct acquisition genotype. After removal of recombinant and hypermutated sequences, the median intra-lineage DNA distance at the first time point was very low: 0.05% for REN (IQR 0.04 and 0.08) and 0.04% for GP (IQR 0.03 – 0.05), with a median inter-lineage DNA distance of 1.68% for REN (IQR 0.94 - 2.66%), and 1.00% for GP (IQR 0.68 – 1.47) **(Supplementary Figure S1)**. This is consistent with diversity observed in acute/early infection (8).

Combined REN and GP sequence analysis identified multilineage infections in 38% of participants, with 37% of infections classified as multilineages by REN alone, and 34% by GP alone (**Table 1**). REN:GP discordant classifications typically involved low abundance secondary lineages (median frequency 0.76%), with 8 of 10 discordant cases having 3 or fewer sequences representing the additional lineage.

### Transmission bottleneck

We investigated if VRC01 treatment either restricted or relaxed the transmission bottleneck, and whether there was a difference between the trials **(Figure 1A and 1C)**. While differences in the VRC01 10mg/kg (29%) and the 30 mg/kg (53%) groups were seen in the Africa trial, the differences were not statistically significant. No difference between the VRC01 groups combined and the placebo group (38%) was observed. In the Americas trial, the frequency of multilineage infection was very similar across all groups, with 41% and 36% of participants in the VRC01 10 mg/kg and 30 mg/kg groups, respectively, and 39% of participants in the placebo group. In addition, there was no difference in the frequency of multilineage infections between the placebo groups in the Africa and Americas trials (38% and 39%, respectively). Lastly, there was no difference between the placebo and VRC01 groups when dose groups were pooled within a study or when groups were pooled across studies.

**Figure 1.**
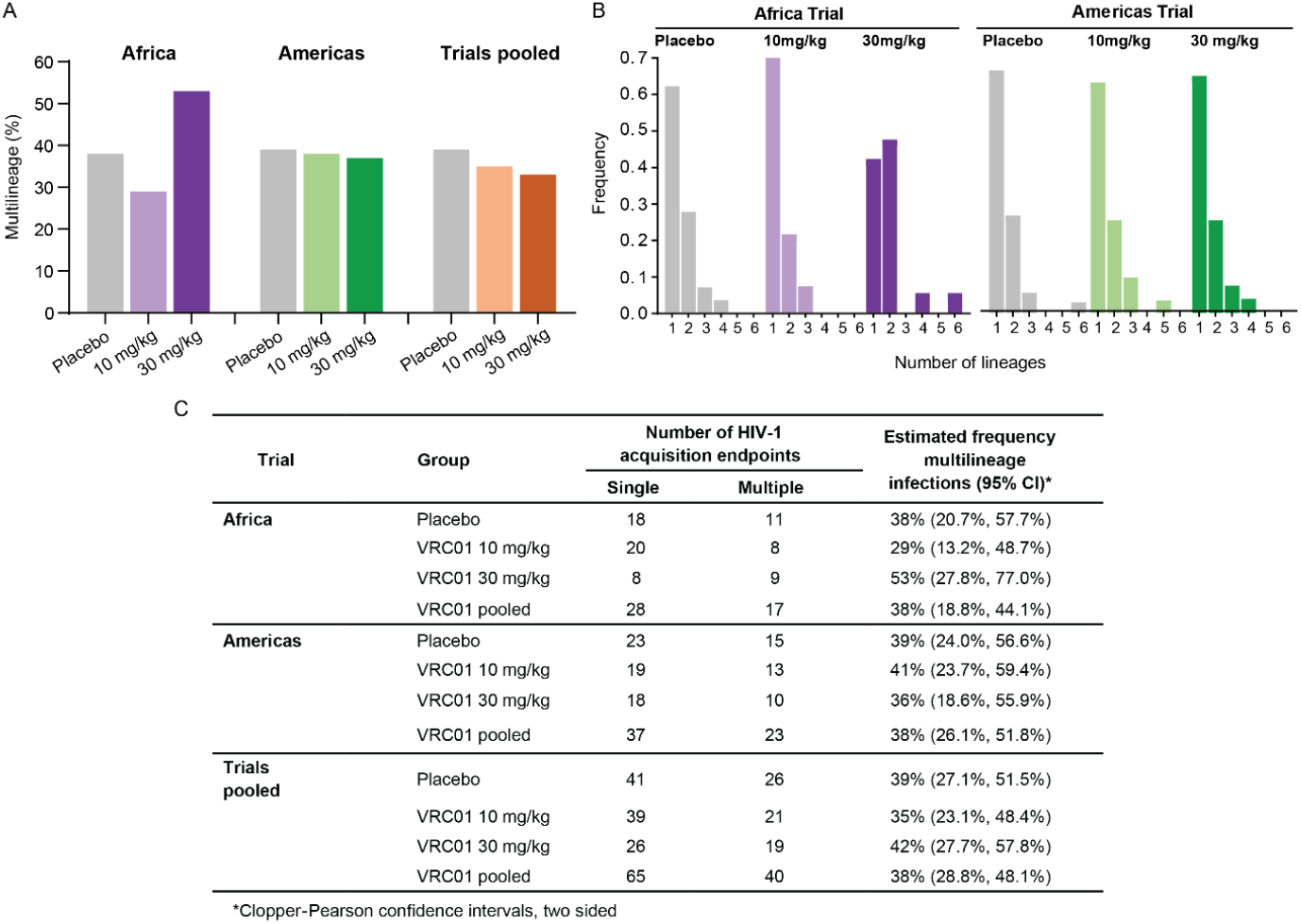
Frequency of multiple transmitted founder lineages (TFL) following HIV diagnosis across and within AMP trials. Gag-Δpol (GP) and rev-env-Δnef (REN) sequences were analyzed to identify individuals with multilineage infection, defined as the presence of more than one lineage in either gene region or at either sampling time point. **A)** Bar graph comparing the percentage of multilineage infections across trial groups: Africa trial (purple); Americas trial (green); placebo grey; pooled trial analysis (orange). **B)** Histogram illustrating the distribution of the number of REN lineages per individual ranging from 1 to 6. **C)** Estimated frequencies of multilineage infections with Clopper-Pearson 95% confidence intervals. No statistical difference was found between groups within trials, or in pooled trial analyses.

Over 90% of individuals acquiring three or fewer lineages. The number of intra-host TFLs was not different between groups within each trial, nor between trials **(Figure 1B)**. The maximum number of lineages detected based on REN sequencing was six, and based on GP sequencing was nine **(Supplementary Figure S2)**.

### Minor viral populations

Minor viral populations, defined here as lineages comprising <5% of the total sequence count, were present in 20% of individuals (35/172 REN; 34/172 GP). Lineages present at a frequency of less than 1% were observed in 12 individuals, for both REN and GP **(Figure 2, Supplementary Figure S2)**. AMP participants had an overall frequency of 38% multilineage infections. If minor lineages in REN were excluded, this number dropped to 22%, similar to the 25% estimated by Baxter et al., (16), demonstrating that increased sequencing depth accounted for the higher-than-expected frequency of multilineage infections. Similarly, following the removal of minor lineages in GP, only 23% of participants would have been determined to have multilineage infections **(Supplementary Figure S2)**.

**Figure 2.**
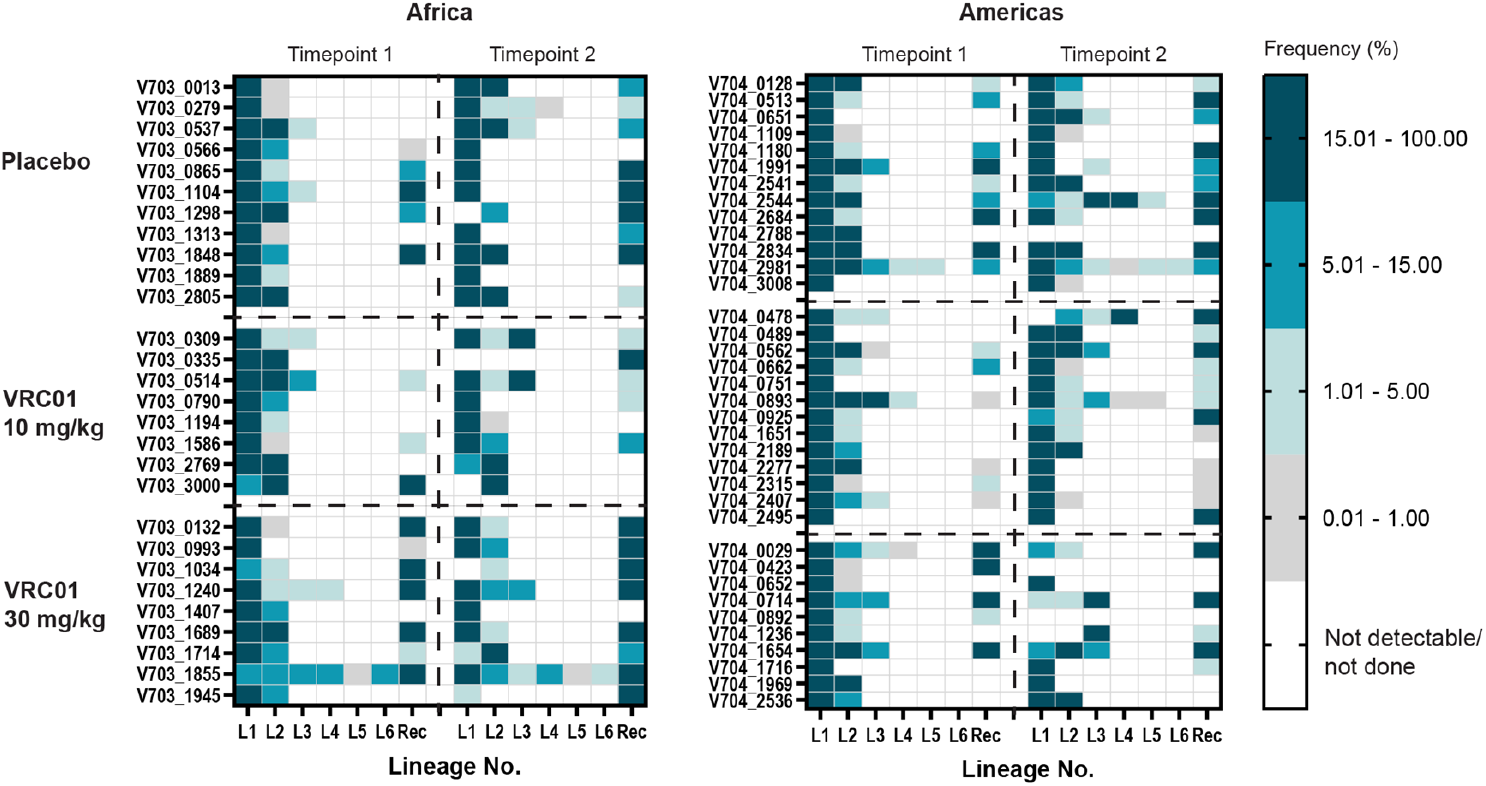
Heat plots illustrating the frequency (%) of rev-env-Δnef (REN) lineages (L1-6) and recombinant sequences (Rec), at two time points in the Africa and Americas trials. Time point 1 corresponds to the approximate time of HIV diagnosis, and time point 2 was sampled on median of 13 days (IQR 6-21 days). Each cell represents the percentage of sequences belonging to a specific lineage or recombinant form, colored from white (not detectable or not done) to dark teal (present in >15% of sequences).

As VRC01 may suppress sensitive viral populations post-acquisition, we assessed whether there was an increased frequency of minor virus populations in the VRC01 arms (pooling across trials and groups). There was no difference in the frequency of minor REN lineages among the individuals in the placebo group (24%; 16/67) compared to the VRC01 group (20%; 21/105). The frequency of minor GP lineages in the placebo group was similar to those in REN (25%; 17/67), although lower in the VRC01 group (14%; 15/105).

Recombination was pervasive, as revealed in individuals with multilineage infections. At the first time point, recombinants were detected in REN in 62% of these individuals in the Africa trial, and 64% in the Americas trial, and 69% and 66% of individuals, respectively, in GP. Pooling across trials and groups, the mean number of unique recombinants generated per day, among all participants for whom recombinants were detected was 0.65 (0.02-7.65) in REN and 0.48 (0.02-2.33) in GP.

### VRC01 discordant phenotypes in participants with HIV multilineage infections

VRC01 neutralization sensitivity was assessed using pseudoviruses generated from representative sequences of each TFL (1, 2). In single-lineage infections, viruses from the pooled VRC01 group (n=65) were significantly more resistant to VRC01 than those from the placebo group (n=41) (Mann-Whitney test: p<0.001) **(Supplementary Figure S3)**.

Of the 66 individuals with multilineage infections, 41 had more than one clone representative of the different lineages and linked to VRC01 neutralization data. One of these participants (V703_1240, VRC01 group) was infected with two HIV strains (dual infection), which may have occurred from two different infecting partners. Of the remaining 40 infections, 40% (n=16) had pseudoviruses with greater than a 3-fold difference in their VRC01 IC80 titres (referred to as VRC01 discordant phenotype) **(Table 2; Figure 3A-F)**. VRC01 discordant infections at diagnosis were more frequent in the treatment group (11 of 24, 46%) compared to placebo (5 of 16, 31%), however this difference was not statistically significant. Notably, three VRC01-treated individuals harbored both highly sensitive (IC80 <1 µg/ml) and highly resistant (IC80 >77 µg/ml) viruses at diagnosis, with differences in sensitivity ranging from 97-to 704-fold (**Table 2**). A trend of increasing intra-host neutralization differences was observed across placebo, low-dose, and high-dose VRC01 groups (Jonckheere-Terpstra test, p=0.072; **Figure 3H**). These findings suggest that an environment around the time of acquisition allowed both VRC01 sensitive and resistant viruses to co-exist.

**TABLE 2:** Maximum-likelihood pairwise DNA distances between and within a lineage of individuals who acquired viruses with VRC01 discordant phenotype (>3 fold difference in IC80) showing fold IC80 difference (*) and inter-lineage DNA distances (**) between the clones and their corresponding lineages with lowest and highest IC80 values.

| Treatment Group | PtID | No. Lineages with IC80 data (total number lineages) | IC80 (µg/ml) |  |  | DNA Distance (mean) |  |
| --- | --- | --- | --- | --- | --- | --- | --- |
|  |  |  | Lowest | Highest | Fold difference* | Intra-lineage | Inter-lineage** |
| Placebo | V703_0537 | 3 (3) | 0.07 | 0.26 | 4 | 0.03% | 1.89% |
|  | V703_2805 | 2 (2) | 1.29 | >100 | 78 | 0.03% | 7.42% |
|  | V704_1109 | 2 (2) | 6.26 | 55.00 | 9 | 0.03% | 1.84% |
|  | V704_2544 | 3 (5) | 2.06 | 14.77 | 7 | 0.02% | 6.50% |
|  | V704_2981 | 4 (6) | 4.93 | 16.42 | 3 | 0.03% | 2.42% |
| VRC01 10 mg/kg | V703_0790 | 2 (2) | 13.02 | >100 | 8 | 0.09% | 0.28% |
|  | V703_2769 | 2 (2) | 0.41 | 77.56 | 189 | 0.05% | 5.47% |
|  | V703_0514 | 3 (3) | 0.14 | >100 | 714 | 0.07% | 6.11% |
|  | V704_0478 | 3 (3) | 10.24 | 37.85 | 4 | 0.03% | 1.30% |
|  | V704_0893 | 3 (5) | 0.81 | 3.58 | 4 | 0.07% | 0.54% |
|  | V704_2315 | 2 (2) | 0.93 | 3.42 | 4 | 0.04% | 1.43% |
| VRC01 30 mg/kg | V703_1714 | 2 (2) | 1.03 | >100 | 97 | 0.14% | 1.25% |
|  | V704_0029 | 2 (4) | 1.06 | 3.72 | 4 | 0.06% | 3.37% |
|  | V704_0892 | 2 (2) | 1.23 | 5.40 | 4 | 0.13% | 1.02% |
|  | V704_1236 | 2 (3) | 2.25 | 6.38 | 3 | 0.03% | 1.83% |
|  | V704_1654 | 2 (3) | 3.14 | >100 | 32 | 0.07% | 1.46% |

**Figure 3.**
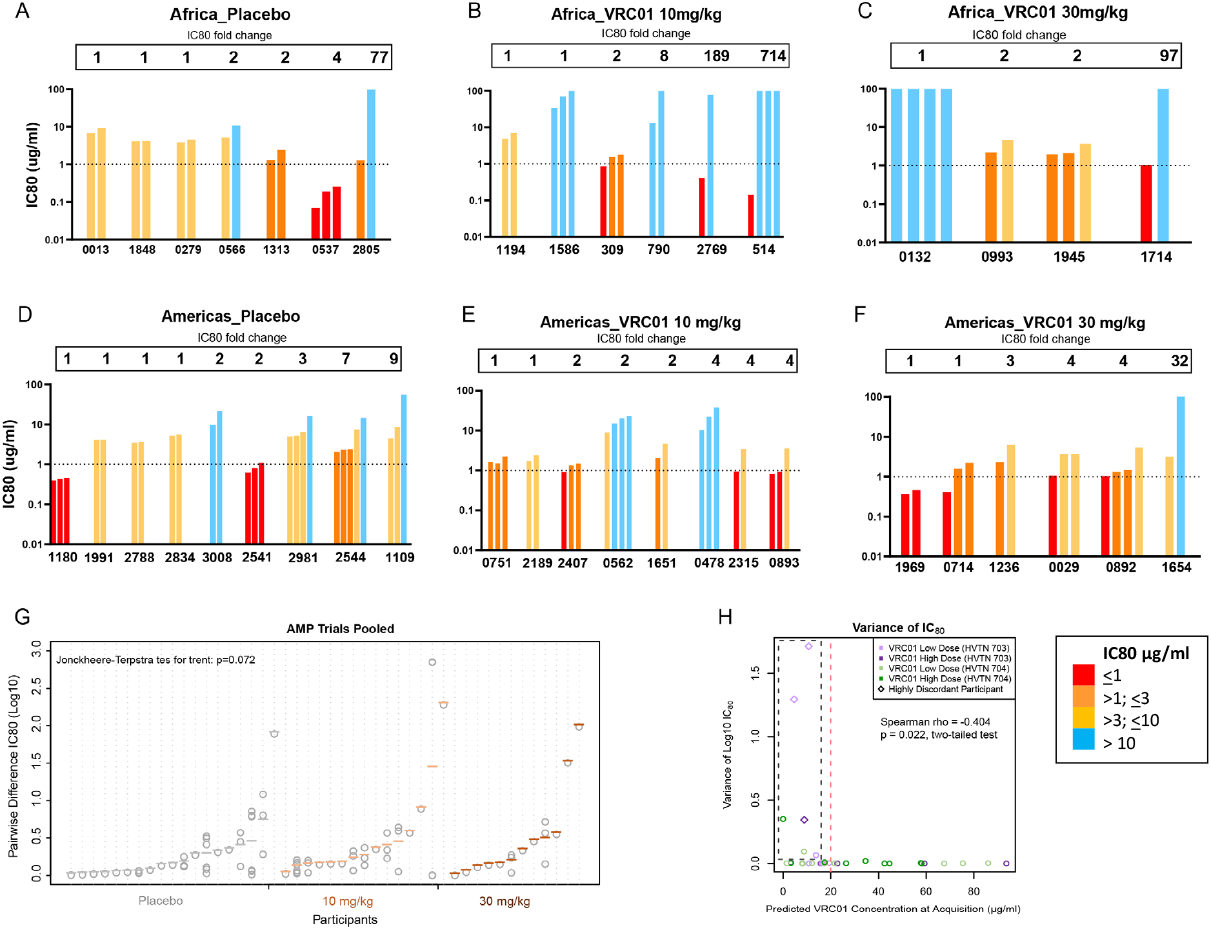
VRC01 discordant phenotypes in AMP participants with multilineage HIV infection. IC80 data shown for participants with neutralization data from multiple clones at the first time point. **A-C)** Africa trial and **D-F)** Americas trial: each bar is one virus from a given. Fold change between the lowest and highest IC80 shown above the bars. **G)** Log IC80 difference between placebo and VRC01 groups (trials pooled). Grey circles show individual pairwise IC80 differences, with colored lines showing the mean pairwise difference placebo (dark grey), VRC01 10 mg/kg (light brown), VRC01 30 mg/kg (burnt orange). Participants with two lineages have one measurement; those with ≥3 have more. Sorted by mean pairwise difference. **H) Variance in IC80 of all sequences (y-axis) vs. predicted VRC01 concentration at acquisition** (estimated date of diagnosable infection minus 7 days). Black dashed box highlights 7 participants with high IC80 variance and low predicted concentration, three of which are slightly above baseline. Above 20 µg/ml predicted concentration (vertical red dashed line), Log10 IC80 values show little variability.

To determine whether a VRC01 resistance phenotype in discordant virus infections was present at acquisition rather than arising through post-acquisition evolution, we compared intra- and inter-lineage sequence differences (**Table 2**). In contrast to the median intra-lineage distances of 0.05% (IQR 0.04 and 0.08); the median an inter-lineage distances of 1.83 were much higher (IQR 1.25-2.70) and generally did not overlap with the intra-lineage values, supporting their classification as distinct infecting lineages (**Supplementary Figure S4**). There was one exception (participant V703_0790) in which the lineages overlapped while differing by only five distinguishing mutations. Thus, we could not exclude the possibility that resistance emerged in this individual after acquisition.

### Influence of VRC01 concentration on co-acquisition of sensitive and resistant phenotypes

We hypothesized that declining VRC01 antibody concentrations between infusions could allow a window of opportunity for co-acquisition of susceptible and resistant viral variants from an infecting partner (18, 19). In this scenario, with relatively weak antibody pressure as antibody levels fell below the protective threshold, resistant viruses would still have a selective advantage while susceptible viruses would escape neutralization. To explore this possibility, the concentration of VRC01 at the estimated time of acquisition was predicted and compared to the variance of IC80s between lineages in the VRC01 groups **(Figure 3G)**. Overall, there was greater intra-participant variance in IC80s at lower predicted VRC01 concentrations (Spearman Rho = -0.404, p=0.022 two-tailed test). In contrast, when the predicted VRC01 concentration was >20 µg/ml, there was limited IC80 variability. The four individuals with the highest variance also had low predicted VRC01 concentrations at the estimated time of acquisition (< 10.73 µg/ml) (V703_0514, V703_1714, V703_2769, V704_1654). Of these, three individuals in the VRC01 treatment groups acquired viruses that were highly sensitive (IC80 < 1 ug/ml) and highly resistant to VRC01 (IC80 >70 ug/ml). The relationship between predicted VRC01 concentrations at estimated acquisition, and the proportion of resistant sequences was also analysed (**Supplementary Figure S5A)**. At an IC80 >3 for resistance we observed a trend of increasing proportion in resistant sequences with increasing predicted concentrations of VRC01 (p=0.054; Spearman Rho, two tailed).

### Viral lineage growth kinetics in participants co-infected with VRC01 sensitivity discordant viruses

Six individuals that acquired virus isolates with an IC80 ≤ 1µg/ml (sensitive) and ≥ 3 µg/ml (resistant) and had sequences from two timepoints **(Figure 4)**. Sequences within a lineage were designated VRC01 sensitive or resistant genotypes based on their genetic relatedness to the pseudovirus evaluated in the neutralization assay. We also screened sequences for evidence of evolution in the VRC01 epitope (20-25).

**Figure 4.**
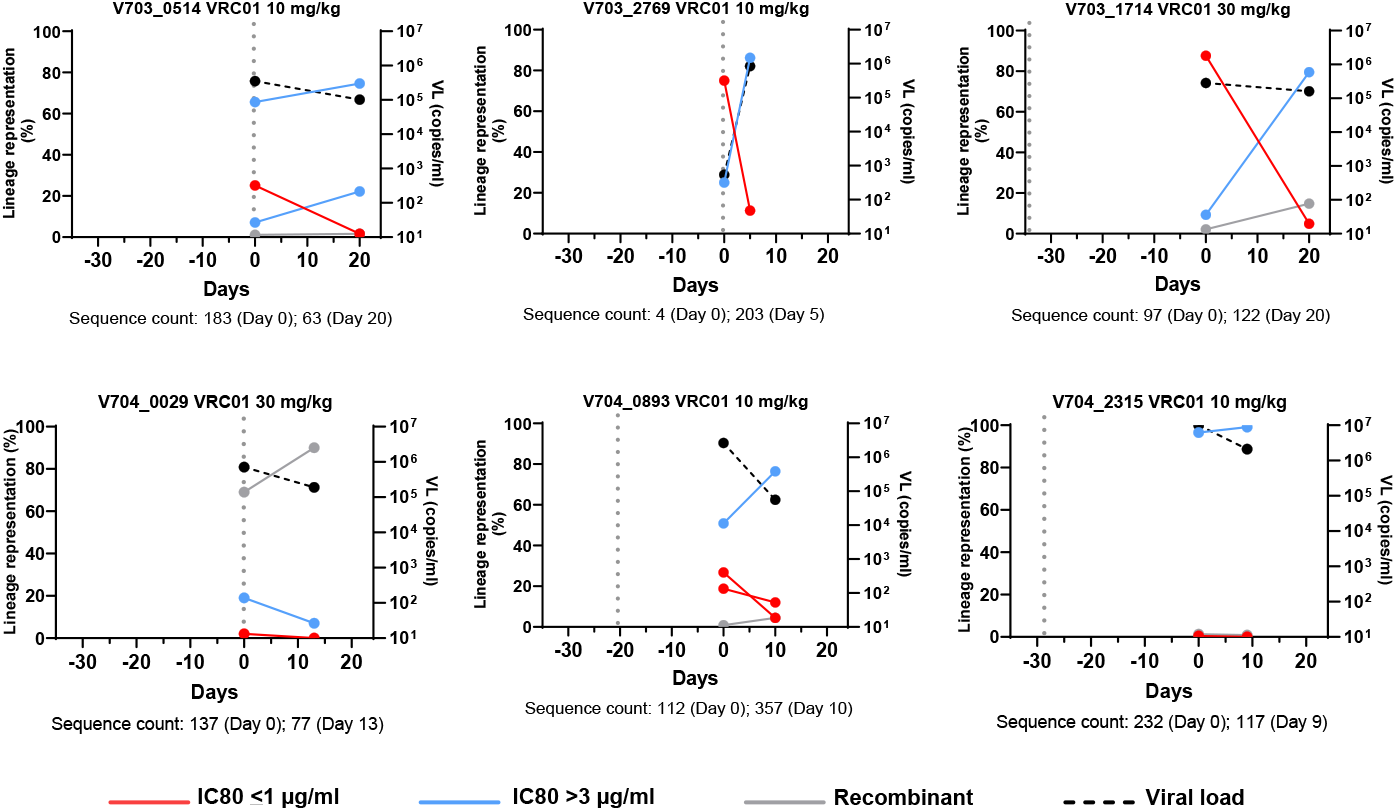
Lineage kinetics in participants who acquired both sensitive (red line) and resistant (blue line) viruses. Lineages were assigned by IC80 of their corresponding transmitted founder pseudovirus. Day 0 represents the first RNA positive timepoint, and the second timepoint is shown in days relative to day 0. Sequence count at each timepoint are indicated below each graph. The sampling interval ranged from 5-20 days. The vertical grey dotted line indicates the timing of the last VRC01 infusion relative to day 0. PSV IC80 values: V703_0514 IC80 >100, >100, and 0.14 µg/ml; V703_2769 IC80 77.56 and 0.41 µg/ml; V703_1714 IC80 >100 and 1.03 µg/ml; V704_2315 IC80 3.42 and 0.93 µg/ml; V704_0029 IC80 3.72, 1.06 and 3.70 µg/ml (recombinant); V704_0893 IC80 3.58, 0.91 and 0.81 µg/ml.

In four of the six participants (V703_0514, V703_1714, V703_2769 and V704_0893), the sensitive lineages were found at reduced frequencies at the later time point (5 to 20 days later) **(Figure 4)** and were found only at very low abundance at both time points in the remaining two participants (V704_0029, V704_2315). In three participants (V703_0514, V703_2769 and V704_0029), the last VRC01 infusion (dotted lines in **Figure 4**) was given on the same day as the first sequencing timepoint, and in the remaining three participants the last infusion was given 25 to 30 days prior to sequencing.

The relative frequency of each infecting lineage was plotted **(Figure 4)** (sensitive red, resistant blue, recombinant gray). In one participant (V704_0029) the lineages were at very low abundance with recombinants accounting for 69% and 90% of the sequences sampled at the first and second time points, respectively. Five recombinant sequences carried the D279A mutation (3% of the total of 190 sequences), all sampled at the second timepoint (recombination p-value <0.001). This mutation is located in the Loop D VRC01 contact site and known to confer VRC01 resistance (21, 24, 26, 27). Toggling at this site was also observed, with D279E, D279H and D279N detected, also carried by recombinant sequences (p-values ranging between 0.01 and 10^−5^). Since the parental lineages contained only D279, it appears the A/E/H/N mutations at position 279 arose after viral acquisition in a recombinant backbone. Once a resistance-associated mutation appeared in V704_0029, it was propagated through recombination (**Supplementary Figure S6**).

Resistance was experimentally defined in two of the six individuals (V703_2769 and V703_1714), allowing for detailed analysis of lineage dynamics (28). Participant V703_2769 was infected with viruses with IC80s of 0.4 µg/ml and 78 µg/ml, and resistance was mapped to changes in the β23-V5 loop **(Figure 5A)**. Participant V703_1714 was infected with viruses with an IC80 of 1.0 µg/ml and >100 µg/ml with partial resistance mapped to the G459D mutation **(Figure 5B**). The abundance of sequences with sensitive mutations declined over five days for V703_2769, and over 20 days for V703_1714, along with a concomitant increase in abundance of sequences with resistance mutations **(Figure 5A and B)**. Recombination was observed in V703_1714 **(Figure 5B and C)** with two inter-lineage recombinants (2% of sequences) at the first time point and 18 (15% of sequences) at the second time point (recombination p-values <0.001). Of these, two early and 14 later recombinants were direct descendants of the two primary lineages, and all inherited the resistance-conferring amino acid D459 - indicating that recombination favoured the resistant genotype (binomial p=0.004), similar to findings in participant V704_0029 described above.

**Figure 5.**
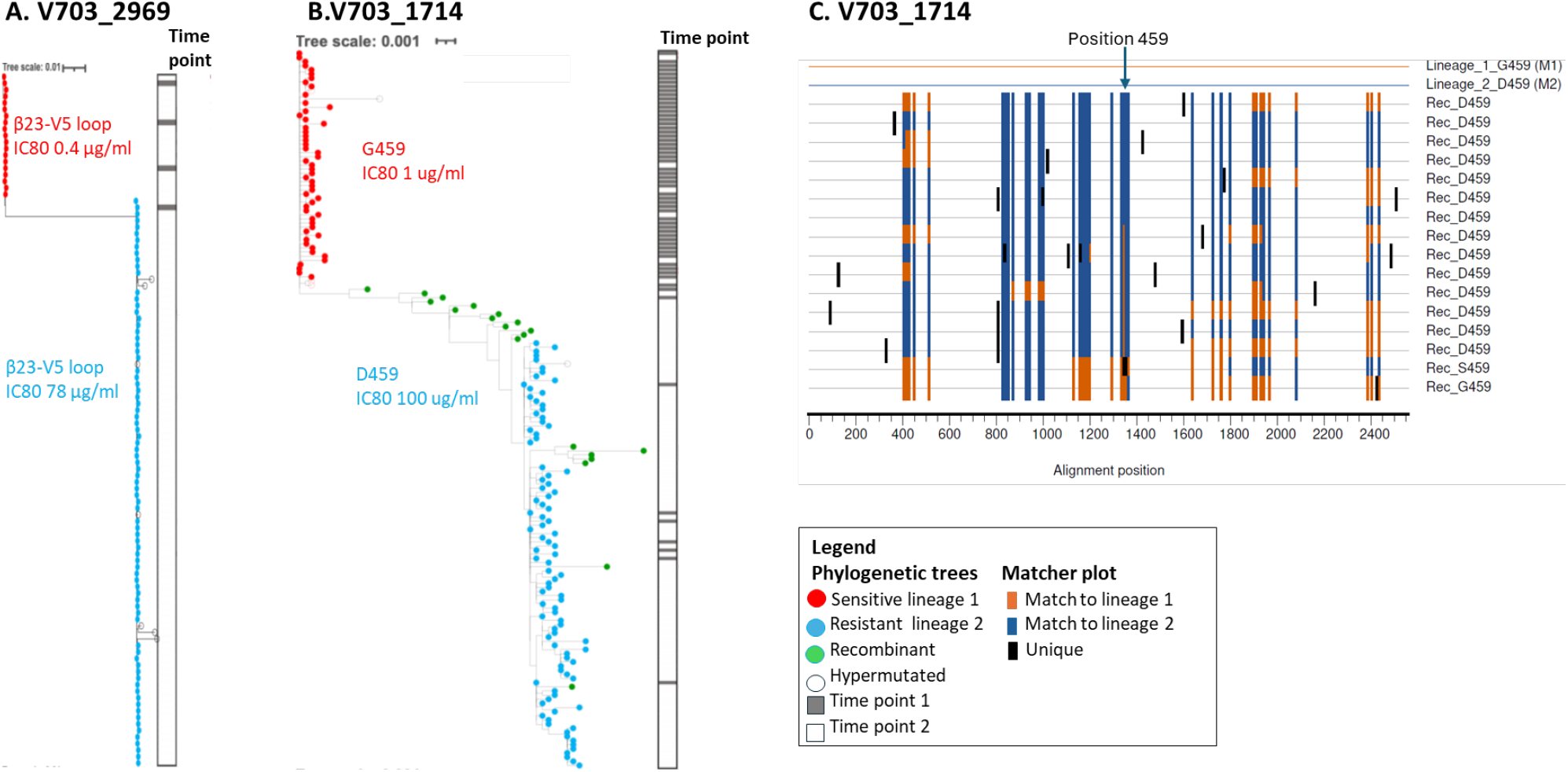
Sequence evolution in two individuals co-infected with VRC01 highly sensitive (IC80 < 1ug/ml) and resistant lineages (IC80>70 ug/ml). In both individuals, VRC01 resistant amino acids were located in the β23-V5 loop (Cohen et al., 2025). Phylogenetic trees of *env* sequences from V703_2769 (A) and V703_1714 (B) and show sequences with sensitive mutations (red dots), resistant mutations (blue dots), and recombinants (green dots). Sequences were obtained from 2 sample time points: 5 days apart for V703_2769, and 20 days apart for V703_1714. The ribbon on the right of each tree illustrated sampling times: grey for time points 1 and white for time point 2. (C) Highlighter plot for V703_1714 illustrating recombination patterns. Each row represents a single env sequence, annotated with either G459 (sensitive) or D459 (resistant) mutations. Vertical tick marks indicate shared site: orange for lineage 1, dark blue for lineage 2, and black for unique changes. The arrow marks the HXB2 position 459

## DISCUSSION

Long-read deep sequencing made it possible to characterise the impact of VRC01 on breakthough infections at a high-resolution. This level of sensitivity was crucial for detecting subtle acquisition and post-acquisition effects that were not identified in previous sieve analysis based on the evaluation of a single sequence per lineage (Juraska et 2024). Using high-resolution deep sequencing (median sequence number 151), multilineage infections were observed at similar frequencies in the placebo group (38%, n=67, pooled) and the VRC01 group (39%, n=105, trials and dose groups pooled). This is a higher rate than reported in a recent meta-analysis, which used lower-sensitivity sequencing approaches (median sequence number 20): founder acquisition probabilities of 30% in men who have sex with men and 21% in women (16). The frequencies observed in AMP were 1.3 to 1.9 times higher and there was no difference between the placebo groups in the two trials (39% in the Americas trial and 38% in the Africa trial). Multilineage infection rates were also similar across geographic regions and circulating subtypes (mainly subtype C in the Africa trial, and subtypes and recombinants B, F and BF in the Americas trial). Interestingly, the number of lineages per participant remained low, with three or fewer lineages detected in 90% of participants, suggesting even with improved sequencing technology, there is still an “acquisition bottleneck”.

Deep sequencing revealed that low frequency viral lineages (< 5% by abundance) were relatively common, detected in ∼20% of individuals in both the placebo and VRC01 groups. These minor lineages contributed to the higher frequency of multilineage acquisitions identified in this study (16). Multilineage infection allows more rapid immune evasion post-acquisition through recombination, which is a more frequent event per viral replication cycle compared to point mutations (29). From a pathogenesis perspective, high viral diversity including multiple founder infections, has been associated with faster disease progression (30-32). This would need to be re-evaluated in light of the prevalence of minor lineages not detectable or considered in previous studies. The reasons for the low abundance of some lineages are unclear, with possible influences including viral fitness, host immunity (innate or possibly adaptive), availability of target cells or stochastic effects during early seeding and population growth.

In single-lineage infections, there was a significant association between VRC01 treatment and IC80 compared to the placebo group. Among individuals with multiple TFL, 40% were infected with viruses that differed in VRC01 sensitivity (IC80 > 3-fold difference).However, the distance between lineages generally exceeded the expected mutation rate for the duration of infection (based on the estimated time of infection and an average mutation rate per nucleotide per generation cycle of 2.16×10-5)(9) (35), and the distances within and between lineage distances did not overlap. This suggests that the discordant phenotype is a property of the transmitted lineage rather than the result of post-acquisition evolution. The coexistence of both VRC01-sensitive and -resistant viruses has also been documented in chronic infection (18).

Higher predicted concentrations at acquisition trended toward selecting resistant viruses (IC80 >3 µg/ml), suggesting it is possible that waning levels of VRC01 between infusions enabled acquisition and outgrowth of these highly sensitive viruses alongside resistant viruses. We hypothesized that there may be a threshold at which VRC01 levels are sufficient to select for resistant viruses, but were too low to fully prevent acquisition of sensitive ones. Indeed, greater variance in IC80 was observed at predicted VRC01 concentrations that were less than 20 µg/ml at acquisition. While statistically significant, this relationship may be stronger than indicated by the Spearman correlations and p-values because the uncertainty in estimating VRC01 levels at acquisition reduces statistical power. These observations underscore the importance of maintaining consistently high monoclonal antibody levels throughout the dosing interval in future prevention strategies.

Six individuals, all in the VRC01 groups, acquired both highly sensitive (IC80 < 1 µg/ml) and resistant (IC80 > 3 µg/ml) viruses. This allowed study of the effect of VRC01 in an environment where viruses with different VRC01 sensitivities replicated under similar conditions. We found a decrease in sensitive viruses over time, suggesting that VRC01 initially suppressed replication of sensitive viral populations post-acquisition. Recombination was pervasive, and in one participant in the treatment group (V703_1714) for which the resistance mutation was mapped (28), with the emerging recombinant viruses were more likely to inherit the VRC01-resistant mutation (p=0.004 binomial).

We assumed that the viral sequence closest to the lineage-specific consensus (‘mindist’) best represented the transmitted founder of that lineage (1, 2). While early changes in the viral genome between acquisition and dissemination cannot be ruled out, there is sufficient data from human transmission studies and animal studies to support this (5, 7, 8, 33). This assumption may be violated if there is a long period between acquisition and sequencing with stochastic effects (34) or post-acquisition selection due to adaptive immune responses (35). In one individual (V704_0029), VRC01 was present at the time of acquisition and sequencing of two closely spaced time points, allowing identification of putative VRC01-induced *de novo* escape. However, this only involved mutations at one position, and was not sufficient to change the lineage classification

This study advances our understanding of how broadly neutralizing antibodies (bNAbs), when used for HIV prevention, exert selective pressure on the virus during the earliest stages of infection. Deep sequencing was essential for identifying these early infection dynamics and subtle resistance patterns. It provided high-resolution insight into how HIV adapts under immune pressure, particularly in the context of multiple transmitted founder lineages. Because resistance can emerge rapidly in these settings, understanding the early viral landscape is crucial. Mapping how HIV evolves under antibody pressure provides a valuable framework for assessing next-generation prevention tools aimed at staying ahead of viral escape. Our data support the use of combination bNAbs with alternative sites of activity to VRC01, similar to the approach used with antiviral therapy. While data are required to determine if post-acquisition selection is reduced with combination therapy, use of the technologies employed in this investigation should be helpful in defining these issues.

## MATERIALS AND METHODS

### Ethical statement

All work described here complied with all relevant ethical regulations. The Institutional Review Boards/Ethic Committees of participating clinical research sites (CRS) approved the studies, which were conducted under the oversight of the NIAID Data Safety Monitoring Board (1). All participants gave written informed consent. The antibody neutralization assays were approved by the Duke University Health System Institutional Review Board (Duke University) through protocol no. Pro00093087 and the University of the Witwatersrand Human Research Ethics Committee (protocol no. M201105). Viral genome sequencing at the University of Cape Town was approved by the UCT Human Research Ethics Committee (HREC reference no. 176/2017) and was considered exempt at the University of Washington.

### Study populations

HVTN 703/HPTN 081, referred to as the Africa trial, enrolled 1,924 women with a high likelihood of HIV acquisition in Botswana, Kenya, Malawi, Mozambique, South Africa, Tanzania, and Zimbabwe. HVTN 704/HPTN 085, referred to as the Americas trial, enrolled 2,704 men who have sex with men with a high likelihood of HIV acquisition from Brazil, Peru, Switzerland, and the United States. Participants were randomized and given infusions of either 10 mg/kg VRC01, 30 mg/kg VRC01, or saline (placebo) (1:1:1 ratio) every eight weeks, with HIV testing scheduled at 4 week intervals (1, 36). Positive HIV RNA diagnosis was confirmed with a second independent sample. The last infusion was scheduled at the week 72 visit, and the primary study endpoints included participants who were diagnosed by the week 80 visit. This report only includes individuals who met the primary endpoints.

### Viral genome sequencing

Sequencing was performed using the Pacific Biosciences single molecule real-time (SMRT) approach with unique molecular identifiers (UMI) as described (17). In brief, during HIV cDNA synthesis, each viral genome was tagged with a UMI, followed by amplification of an estimated 100 or more templates corresponding to the *gag*-Δ*pol* (GP) genomic region (2.5kb; HXB2 positions 790 - 2292) and the *rev-env-*Δ*nef* (REN) region (3 kb; HXB2 positions 5970 – 9012). Amplicons were sequenced on the PacBio platform using Sequel and Sequel II instruments. Circular consensus sequences (CCS) were generated, and sequence data processed using a custom bioinformatics pipeline, PORPIDpipeline (http://github.com/MurrellGroup/PORPIDpipeline) (17). Nucleotide and amino acid alignments were generated using Muscle5 (37) and refined using Geneious (https://www.geneious.com). Identical sequences were collapsed for manual alignment review, and three rounds of independent review were performed. Nucleotide diversity was calculated using Phylogenetic estimation using Maximum Likelihood (PhyML) (https://github.com/stephaneguindon/phyml) with the General Time Reversible (GTR) model of evolution in DIVEIN (38). Participants in the Africa trial predominantly acquired env subtype C (98%), with one G and one A1C recombinant identified (2, 39). Participants in the Americas trial predominantly acquired env subtype B (73%), subtype F (11 %), and recombinant B/F (11%), with subtype C and D, and recombinant forms A1/B, B/D and CRF47_BF also detected (2, 39).

### Lineage determination

These methods are described in detail by Mullins et al. (in prep). In brief, to identify individuals with multilineage acquisitions, maximum-likelihood phylogenetic trees were generated. Nucleotide changes from the most common sequence at the first time point were visualized using highlighter and matcher plots in Phylobook (https://github.com/MullinsLab/phylobook) (40). Lineage assignment was supported by k-medoid clustering, the Poisson Fitter tool (33) and gap-methods (41). Hypermutated sequences were identified using the LANL Hypermut tool (https://www.hiv.lanl.gov/HYPERMUT) and excluded from lineage analysis. The minimum number of nucleotide changes to define a lineage for both GP and REN was four, with a minimum of two shared changes across a lineage used to define a recombinant. Maximum likelihood pairwise distance distributions were generated using PhyML v3.3 (42) within DIVEIN (38).

Visual inspection using Phylobook (40) and the Recombination Analysis Program (RAPR, https://www.hiv.lanl.gov/RAP2017) were used to screen for recombinants (the latter using a significance threshold of FDR q=0.2) (43). To calculate recombination rates, statistical support for each unique recombination event was obtained using the Wald–Wolfowitz Runs Test (43) with the most common sequence in each distinct lineage as a potential parental strain. In participants with VRC01 discordant phenotypes, RAPR was used to estimate how recombination affected the prevalence of the escape/resistance mutations. This included sequences from all available time points and a multiple testing FDR threshold of q<0.2.

### Pseudovirus production and neutralization assays

The *rev-env* portion of sequences representative of the different viral lineages were synthesized and cloned, and HIV Env-pseudotyped viruses generated as described (1). 293T/17 cells were co-transfected with *env* plasmids and an *env*-deleted backbone vector (pSG3delEnv). Pseudoviruses were then titrated to determine the optimal dilutions to use in the TZM-bl neutralization assay (44, 45). Neutralization was measured as a function of reduction in luciferase reporter gene expression (due to the presence of bNAbs) after a single round of HIV Env-pseudotyped virus infection of TZM-bl target cells. Neutralization titres are expressed as the concentration of VRC01 at which relative luminescence units (RLU) were reduced by 80% (IC80) relative to virus control wells after subtraction of background RLU in cell control wells.

### VRC01 discordant phenotypes

Intra-participant differences in IC80 were calculated as the fold change between the most sensitive and most resistant clones. Discordant phenotypes were defined when two clones were greater than 3-fold different in their IC80 values, a threshold which considers assay variability (45). Clones classified as recombinant were excluded from this analysis. The calculation of estimated dates of diagnosable acquisition (EDDA) is described in detail by (19). In summary, the EDDA was estimated using a Bayesian posterior distribution, which combined independent inputs from diagnostic timing estimators with combined gag-Δpol and *rev-env-Δnef* based estimators utilizing Poisson Fitter 2.0 (46). When two sequenced regions yielded distinct timing estimates (that is, the 95% posterior credible intervals did not overlap), the *gag-Δpol* region was chosen as the final time estimate as this region does not encode a protein targeted by VRC01. In this study, the date of acquisition was assumed to be 7 days prior to the EDDA to account for the infection eclipse phase. VRC01 serum concentrations at acquisition were predicted by a 2-compartment population pharmacokinetics (popPK) model (47, 48). To examine the relationship between VRC01 levels at acquisition and the probability of multiple TFLs, we correlated available IC80 values at the first sequencing timepoint with the participant’s predicted VRC01 concentration at the estimated time of acquisition.

### Statistical analysis

Ninety-five percent confidence intervals (CI) for the frequency of multilineage compared to single lineage acquisitions were estimated by Clopper-Pearson CI (also known as binomial proportion CI, two-tailed p-values). Distributions of IC80 values were evaluated for each primary endpoint case (placebo vs VRC01 dose groups and trials pooled) using the Mann-Whitney U test (two-tailed). Intra-participant differences in VRC01 sensitivity were summarized by the mean of their pairwise-clone log10-transformed IC80 values. The individual mean values were then compared across groups and the three ordered treatment groups placebo, VRC01 10 mg/kg, VRC01 30 mg/kg using the Jonckheere-Terpstra test for trend (one-sided) (49). All statistical tests examining the relationship between VRC01 levels at estimated acquisition and the probability of multiple TFL, including the analyses of intra-participant IC80 variance, and the recombination statistical tests (Runs Test and Binomial Tests) were done with correlation tests using the Spearman Rho coefficient (two-tailed p-values).

Data, Material, and Software availability. The neutralization data underlying the findings of this manuscript are publicly available at the public-facing HVTN website (https://atlas.scharp.org/project/HVTNPublicData/). All individual participant data have been deidentified. All final GP and REN sequences were deposited in GenBank with Accession numbers (XXX-YYY). The GenBank accession numbers for the HIV Env clones used in the TZM-bl target cell neutralization assay are: HVTN 703/HPTN 081 sequences, ON890939–ON891092; HVTN 704/HPTN 085 sequences, ON980814–ON980967.

## ACKNOWLEDGEMENTS AND FUNDING SOURCES

We thank the participants of the HVTN703/HPTN081 and HVTN704/HPTN085 clinical trials; the clinical site staff; the protocol development and study implementation teams; NIH/NIAID Vaccine Research Center for the clinical development and manufacturing of the study product.This work was supported by the National Institutes of Health https://www.niaid.nih.gov/: UM1 AI068614 to LC at HVTN, FHCC; UM1 AI068635 to PBG, YH, HJ at HVTN, SDMC, FHCC; UM1 AI068618 to MJM at HVTN, FHCC; UM1 AI068619 to MSC at HPTN; UM1 AI068613 to MSC at HPTN; UM1 AI068617 to MSC at HPTN; R01 AI152115 to CW, PBG PLM, and LM. PLM and CW and their teams are supported by the South African Medical Research Council Strategic Health Innovations Department. PLM is supported by the South African Research Chairs Initiative of the Department of Science and Innovation and the National Research Foundation (grant no. 98341). DBR is funded by K25 AI155224 and R01 AI186721-01. This study was supported in part by the Bill & Melinda Gates Foundation (CAVD; grant 1032144 to DM; and INV-016189 to JIM. BM was supported, in part, by the Swedish Research Council (2018-02381) and the NIH NIAID (R01 AI157854 - subaward).

## AUTHORS CONTRIBUTIONS

CW, JIM, PBG, L.Corey, MC designed the research; JIM, JHW, CW, CM, CAM, EEG, MR, RR, TB, BM, JH, DHW developed or designed the methodology; DHW, NNM, LC, HZ, TY, AG-N, NN, RT, PC, BL, HK, SB performed the research assays; STK, JAH, LM, DM, SI, NM, MJE, MSC, L.Corey, PGG, LB, MR, REB provided study materials, reagents, computer resources or other analysis tools; CW, JIM, CM, CAM, EEG, TB, BM; DY, WD, RR, AP, SE, NM, MJM, LM, PLM, MSC, LZ, DBR, BMCAM, AdeC, JL, AY, AL, BM, REB, PTE, NB, DM, WD performed data curation or analysis; JIM, CW, CM, CAM, EEG wrote the paper; all authors reviewed and edited the paper.

## Supplementary Figures

Figure S1. REN and GP DNA distances. Intra-lineage and inter-lineage DNA distances at the first sequencing time point only for all participants who met study endpoint. Participants included: V703_REN (n=74), V703_GP (n=63), V704_REN (n=97) and V704_GP (n=96). The upper and lower horizontal lines indicate the inter-quartile range and the middle horizontal line indicates the median.

Figure S2. Heat plots illustrating the lineage frequency (%) of A) rev-env-Δnef (REN) lineages (L1-6) and recombinant sequences (Rec), and B) Gag-Δpol (GP) (L1-9) at two time points in the Africa and Americas trials. Time point 1 corresponds to the approximate time of HIV diagnosis, and time point 2 was sampled a median of 13 (IQR 6-21 days) later. Each cell represents the percentage of sequences belonging to a specific lineage or recombinant form, colored from white (not detectable or not done) to dark teal (present in >15% of sequences).

Figure S3. Distribution of IC80 for primary endpoint cases (trials and treatment groups pooled) single lineage infections. Placebo group shown as grey dots, and the VRC01 group as brown dots. The top and bottom horizontal lines indicated the inter-quartile range, and the bold middle line indicates the median. Median IC80 of the placebo group (n=39) was 1.83 µg/ml and the VRC01 group was 4.87 µg/ml (n=62). Columns compared using Mann-Whitney test, two tailed, *** p-value < 0.0005.

Figure S4. Plots of 16 VRC01 discordant multilineage infections illustrating maximum-likelihood pairwise DNA distances within and between lineages. The medians are indicated by black horizontal lines.

Figure S5. A) Proportion of resistant sequences at IC80 > 3. The plot shows the proportion of resistant sequences within each participant (IC80 > 3 as threshold for resistance) on the y-axis against predicted VRC01 concentration at acquisition. B) Table showing IC80 variance for the four individuals with the greatest IC80 variability (i.e., a larger number of both high and lower values) in participants with a lower predicted VRC01 concentration (< 20 µg/ml). The table shows the number of lineages assayed (from the total lineages), plus the total number of sequences representing each lineage

Figure S6: Nine recombinant sequences found in V7004_0029, each carrying a different escape mutation at position 279. Each panel shows recombinant sequence(s) with the two parental strains at the top of each graph and the mutations colored blue or orange according to the parental strain they matched. (A) Five recombinant sequences, all sharing the same parental strains, namely, the most common sequence in the first founder lineage (TFL1) and the most common sequence found in the second founder lineage (TFL2). Both parental lineages carried the 279D amino acid, hence each of the 5 recombinants acquired the escape mutation de novo or inherited from a prior parental strain that acquired it following HIV acquisition. Escape mutations are color coded to illustrate the different amino acid escape variants. (B-E) Four additional recombinant sequence variants in which parental strains carried one sensitive and one resistance mutation, and the recombinant sequences all inherited the resistant mutation. Parental strains are distinct from the founder lineage strains in panel A and indicated as P1 and P2. The second parental lineage is a recombinant itself, and hence these are all second-generation recombination events. P-values on the left of the graph are from the runs test.

