## Supplemental Material for "Influence of the broadly neutralizing antibody VRC01 on HIV breakthrough virus populations in antibody-mediated prevention trials"

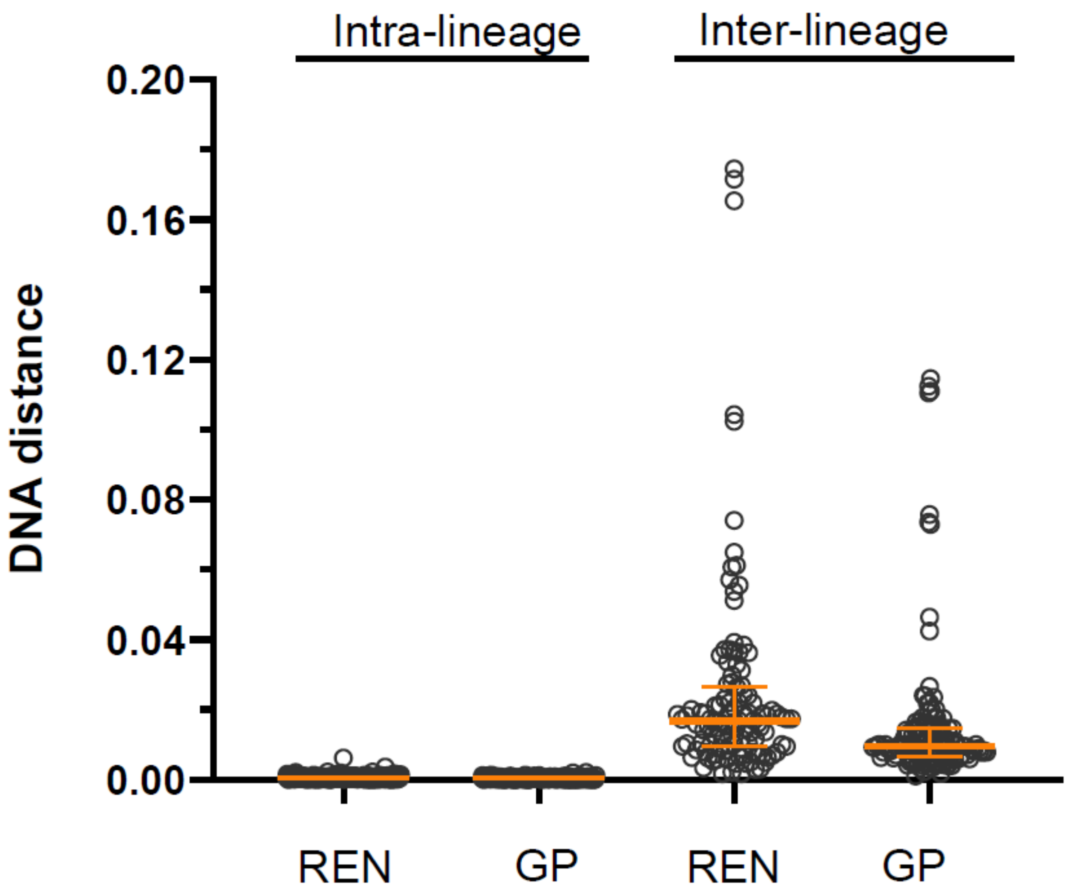


**Supplementary Figure S1. REN and GP DNA distances.** Intra-lineage and inter-lineage DNA distances at the first sequencing timepoint only (i.e. timepoint 1) for all participants who met study endpoint. Participants included, V703_REN (n=74), V703_GP (n=63), V704_REN (n=97) and V704_GP (n=96).

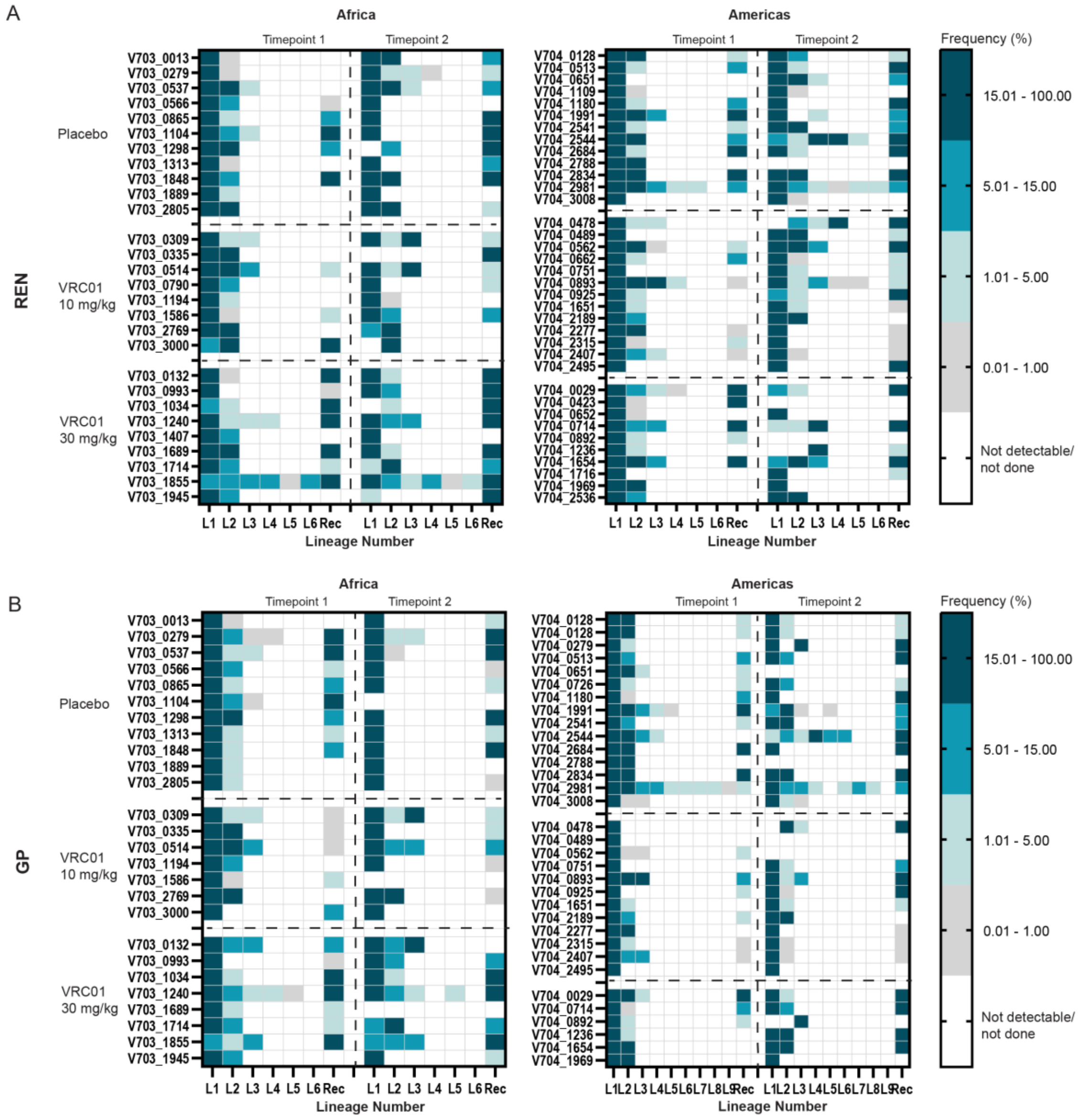


**Figure S2.** Heat plots illustrating the frequency (%) of **A)** rev-env-Δnef (REN) lineages (L1-6) and recombinant sequences (Rec), and **B)** Gag-Δpol (GP) (L1-9) at two time points in the Africa and Americas trials. Time point 1 corresponds to the approximate time time of HIV diagnosis, and time point 2 was sampled on average of 17 days later (IQR 8-20 days). Each cell represents the percentage of sequences belonging to a specific lineage or recombinant form, colored from white (not detectable or not done) to dark teal (present in >15% of sequences).

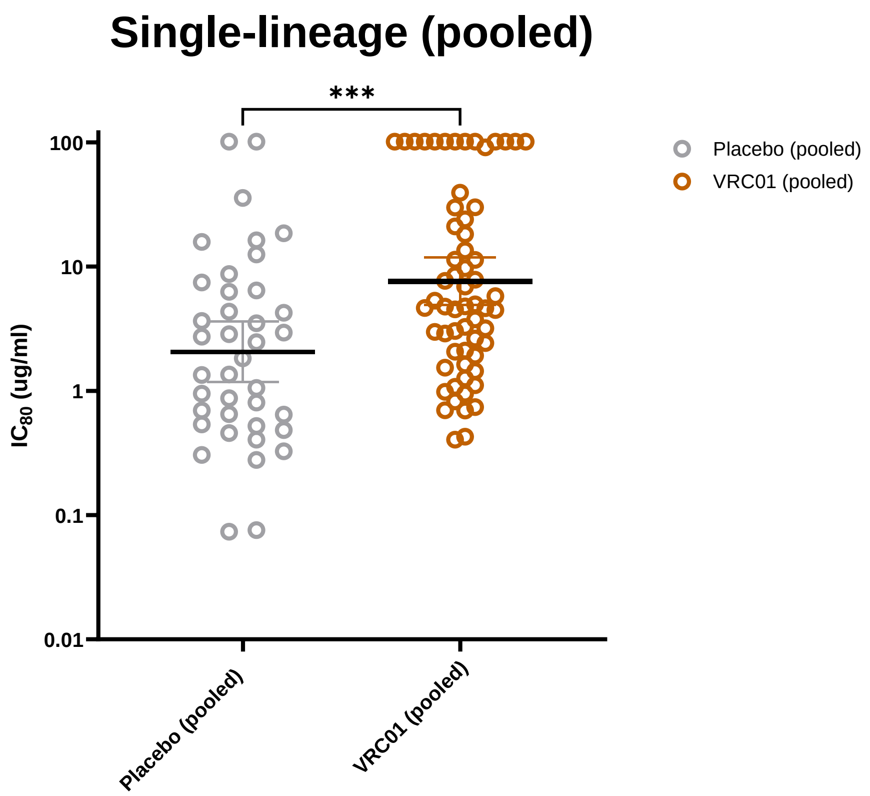


**Figure S3. Distribution of IC80 for primary endpoint cases (trials and treatment groups pooled) single lineage infections.** Placebo group shown as grey dots, and the VRC01 group as brown dots. The top and bottom dotted lines indicated the inter-quartile range, and the dashed middle line indicates the median. Median IC80 of the placebo group (n=39) was 1.83 μg/ml and the VRC01 group was 4.87 μg/ml (n=62). Columns compared using Mann-Whitney test, two tailed, *** p-value < 0.0005.

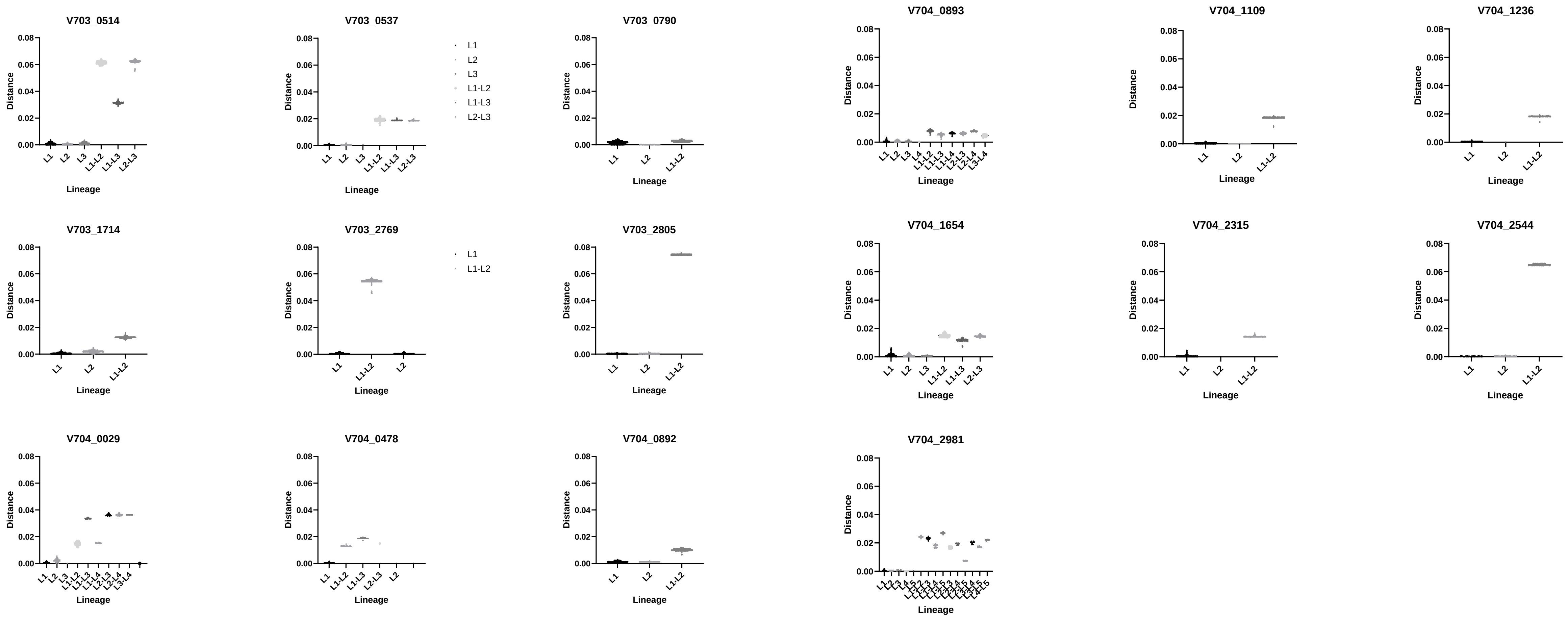

**Figure S4.** Plots of 16 VRC01 discordant multilineage infections illustrating maximum-likelihood pairwise DNA distances within and between lineages. The medians are indicated by black lines.

A.

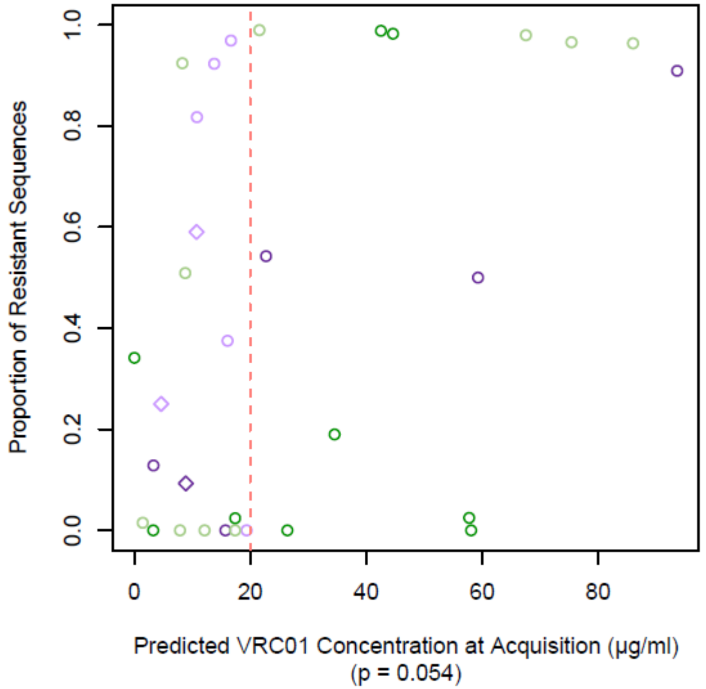


B.

| Log10 variance IC80 | Trial_Ptid_dose | Sequences assayed (from total lineages) | Sequences from sensitive lineage | Sequences from resistant lineage | Sequences not represented |
|---|---|---|---|---|---|
| 1.72 | V703_0514_10 mg/kg | 4 (3) | 46 (25%) | 108 (59%) | 29 (16%) |
| 1.30 | V703_2769_10 mg/kg | 2 (2) | 3 (75%) | 1 (25%) | all lineages represented |
| 0.35 | V704_1654_30 mg/kg | 2 (3) | 8 (6%)* | 35 (28% | 83 (66%) |
| 0.34 | V703_1714_30 mg/kg | 2 (2) | 9 (9%) | 85 (88%) | 3 (3%) |

Ptid = participant id. *IC80 3.1 μg/ml.

**S5. A) Proportion of resistant sequences at IC80 > 3.** The plot shows the proportion of resistant sequences within each participant (IC80 > 3 as threshold for resistance) on the y-axis against predicted VRC01 concentration at acquisition. **B) Table showing IC80 variance for the four individuals with the greatest IC80 variability (i.e., a larger number of both high and lower values) in participants with a lower predicted VRC01 concentration (< 20 μg/ml).** The table shows the number of lineages assayed (from the total lineages), plus the total number of sequences representing each lineage.

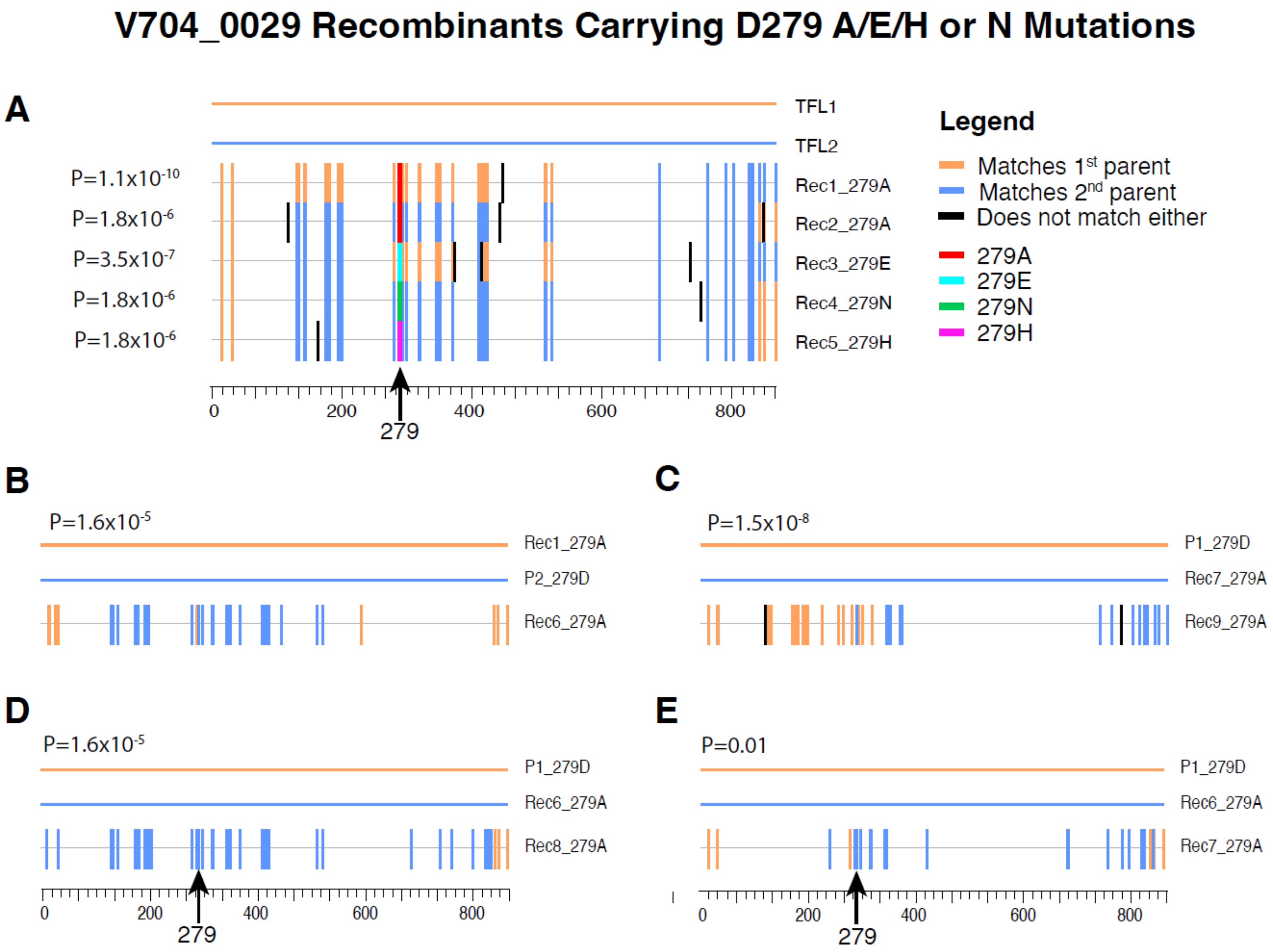


**Figure S6**: Recombination analysis of V704_0029 illustrating preferential inheritance of 279A mutation. Each panel shows recombinant sequence(s) with two parental strains at the top of the graph and the mutations color coded blue or orange according to the parental strain the match. (A) Five recombinant sequences who developed the resistant mutation de novo. B-E four recombinant sequences whose parental strains carried one the sensitive mutation and one the resistant mutaions and the recombinant sequences all inherited the resistant mutation. P-values on the left of the graph are from the runs test.
